# Selective Cancellation of Reactive and Anticipated Movement: Differences in Speed of Action Reprogramming, but not Stopping

**DOI:** 10.1101/2023.10.19.563181

**Authors:** Simon Weber, Sauro E. Salomoni, Mark R. Hinder

## Abstract

The ability to inhibit movements is an essential component of a healthy executive control system. Two distinct but commonly used tasks to assess motor inhibition are the stop signal task (SST) and the anticipated response inhibition (ARI) task. The SST and ARI tasks are similar in that they both require cancelation of a prepotent movement; however, the SST involves cancelation of a speeded reaction to a temporally unpredictable signal, while the ARI task involves cancelation of an anticipated response that the participant has prepared to enact at a wholly predictable time.

33 participants (mean age = 33.3 years, range = 18-55 years) completed variants of the SST and ARI task. In each task, the majority of trials required bimanual button presses, while on a subset of trials a stop signal indicated that one of the presses should be cancelled (i.e., motor selective inhibition). Additional variants of the tasks also included trials featuring signals which were to be ignored, allowing for insights into the attentional component of the inhibitory response. Electromyographic (EMG) recordings allowed detailed comparison of the characteristics of voluntary action and cancellation.

The speed of the inhibitory process was not influenced by whether the enacted movement was reactive (SST) or anticipated (ARI task). However, the ongoing (non-cancelled) component of anticipated movements was more efficient than reactive movements, as a result of faster action reprogramming (i.e., faster ongoing actions following successful selective inhibition). Older age was associated with both slower inhibition and slower action reprogramming across all reactive and anticipated tasks.

## 1. Introduction

### 1.1. Stop signal tasks

Response inhibition refers to the ability to cancel planned or initiated actions. The most commonly used task to assess response inhibition in the laboratory setting is the stop signal task (SST; Leunissen et al., 2017; Logan & Cowan, 1984; Verbruggen et al., 2019). Several variants of the SST exist, though they typically involve a predominance of “go trials” in which the participant must rapidly enact a prescribed movement (e.g., press a button) in response to a “go signal”. Randomly interspersed amongst these go trials are a small proportion of “stop trials” in which the go signal is followed by a “stop signal” which indicates that the participant must cancel their initiated response. This paradigm has been used broadly in recent decades for clinical and non-clinical research in psychology, psychiatry, neuroscience and biology (Verbruggen et al., 2019).

### 1.2. Anticipated response inhibition tasks

An alternative to the SST is the anticipated response inhibition (ARI) task (Coxon et al., 2006; He et al., 2022; Slater-Hammel, 1960). Like the SST, the ARI task is comprised of both go trials and stop trials. However, in a go trial, the participant must enact a response when a moving indicator intersects a stationary target. Typically, this moving indicator is a vertical bar “filling” (i.e., changing colour) from the base upwards, at a constant and predictable rate. The moment that this rising colour change intersects a horizontal target line is when the participant must enact their movement. Accordingly, they predict (or anticipate) when to initiate their response such that the behavioural manifestation (i.e., the button press) occurs to stop the moving indicator at the target. On stop trials, the stop signal occurs *prior* to the point where the moving indicator intersects the target, and the participant must attempt to cancel their prepared response. Typically, this stop signal is represented by the filling bar stopping its upward progress before it reaches the target line.

The SST and ARI tasks are similar in that they both involve the cancelation of a prepotent movement. The key distinguishing feature, however, is that the SST involves cancelation of a reactive response to an unpredictable go signal appearing, whereas the ARI task involves cancelation of an anticipated response that the participant has prepared to enact at a foreseen (i.e., predictable) time. To our knowledge, direct comparisons between these tasks have only assessed behavioural data (He et al., 2022; Leunissen et al., 2017; Wadsley et al., 2022), and it is therefore unknown whether inhibitory network recruitment differs between SSTs and ARI tasks. Given that reactive and anticipated movements differ in their recruitment and modulation of the motor cortex (Bastian et al., 2003; Davranche et al., 2007), it is plausible that the neural mediators of inhibition also differ between these types of movement.

### 1.3. Using electromyography (EMG) to characterise action cancellation

Conventional measures of reactive inhibition for both the ARI task and the SST involve deriving an estimate of stop signal reaction time (SSRT) based on behavioural data. SSRT calculation is based on the ‘horse race model’ of action inhibition (Logan & Cowan, 1984). This model proposes that the outcome of a response following a stop signal is determined by which of two independent processes is the first to reach threshold: a go process (triggered by the go signal) and a stop process (triggered by the stop signal). Conventional SSRT calculations have been called into question based on violations of assumptions in the model underlying them (Bissett et al., 2019; Gulberti et al., 2014; Matzke et al., 2017; Verbruggen & Logan, 2015), with further issues highlighted in regard to using standard SSRT estimations in the ARI task (Matzke et al., 2021). An alternate method of indexing stopping latency involves directly recording muscle activity using electromyography (EMG) while a participant completes a behavioural task involving action cancellation. In successful stop trials, no behavioural response is recorded from the stopped effector (that is, the prepotent movement has been successfully inhibited prior to an overt button press). However, the participant may generate sub-threshold muscle activation, whereby they initiated a motor response, but this was cancelled before enough force was generated to press the button (known as a partial EMG burst). By calculating the delay between the presentation of the stop signal and the peak amplitude of the partial burst (i.e., when covert muscle activation begins to decrease), inhibition latency can be indexed on a single trial level – a measure previously referred to as CancelTime (Jana et al., 2020; Raud et al., 2022; Raud & Huster, 2017). Furthermore, assessing inhibition as it occurs in the muscle removes the ballistic stage of action cancelation (i.e., electromechanical delays between muscle contraction, force generation and registration of a button press), which is subject to variations in experimental conditions, such as the effector with which the participants execute the movement, button stiffness (Gronau et al., 2023), or whether the response requires pressing or releasing a button (e.g., Coxon et al. 2007).

While a number of studies have assessed CancelTime resulting from SSTs, to our knowledge no research currently exists which calculates this on data from an ARI task (though we note that partial burst frequency and amplitude are reported in response to a unimanual ARI task by Coxon et al., 2006). Here, we seek to make direct intra-individual comparisons between CancelTime distributions in both ARI tasks and SSTs. Past research has found SSRT to be shorter in ARI tasks compared to SSTs (Leunissen et al., 2017), however, this finding is not universally observed (Hall et al., 2022; Wadsley et al., 2023), and the calculation of SSRT in these contexts has been rendered questionable (Matzke et al., 2021). Given the lack of previous research, we form no specific hypothesis regarding how CancelTime will differ between these tasks.

### 1.4. Motor selective stopping tasks

Motor selective stopping refers to instances in which we inhibit only a subset of the movement or effectors engaged in an action, while continuing with others, e.g., stopping one hand during a bimanual movement while continuing with the other hand (Aron & Verbruggen, 2008; Coxon et al., 2007; Wadsley et al., 2022; Weber et al, 2023). Motor selective stopping paradigms use selective stop signals that specify which component of a multi-component action should be cancelled. A large body of research demonstrates that performance in motor selective stopping is “sub-optimal”, that is, the reaction time of the continued action component is delayed relative to reaction times in trials where no stop signal was presented (Cai et al., 2011; Claffey et al., 2010; Coxon et al., 2007; Drummond et al., 2018; Majid et al., 2012; Raud & Huster, 2017). While this observation has historically been called the “stopping interference” (SI) effect, Gronau and colleauges (2023) refer to the longer reaction time as a *stopping delay*, reflective of the delayed start time of the new (selected) response, rather than any interference between the inhibition and action mechanisms. This delay indicates that the nervous system does not truly enact selective stopping, but instead facilitates a global inhibitory response (affecting any ongoing movements) and the initiation of a new unimanual response in the non-signalled effector (MacDonald et al., 2014, 2017; Gronau et al., 2023).

Recent work has indicated these RT delays are greater SSTs than ARI tasks (Wadsley et al., 2022). However, it is currently unknown whether this is a result of faster suppression of the bimanual movement (prior to reprogramming of the continued effector), or faster action reprogramming (a shorter time between the suppression of the initial movement and the initiation of the required selective movement). Here, we seek to use EMG derived indices of when activation patterns change at the level of the muscle to add further insights into the underlying processes of motor selective stopping in both SSTs and ARI tasks.

### 1.5. Stimulus selective stopping tasks

A growing body of evidence suggests that inhibitory responses occur to unexpected stimuli irrespective of the behavioural imperative that is associated with them (Wessel & Aron, 2017; Weber et al., 2023). This broadly applied inhibitory response has been theorised to rely on the hyperdirect cortico-basal-ganglia pathway (Chen et al., 2020) and to be distinct from the voluntary cancellation of an action, mediated by the slower indirect pathway (Diesburg & Wessel, 2021). In SSTs and ARI tasks, the infrequent occurrence of stop trials causes both voluntary and involuntary sources of inhibition to become conflated when the stop signal appears. One potential means of teasing these processes apart has been to implement “stimulus selective” stopping. This involves using two perceptually distinct types of infrequent signal during an stopping task, only one of which is associated with the imperative to stop (van de Laar et al., 2011). The other signal (referred to here as an “ignore” signal), bears no behavioural imperative; the participant must simply ignore the unexpected signal and enact their response as though no additional signal has appeared. When examining behavioural results, the extent of the reaction time slowing that occurs in correct stop trials is greater than that which occurs in ignore trials in both SSTs (Ko & Miller, 2013) and ARI tasks (Wadsley et al., 2023). These findings suggest that the attentional component of inhibition does not entirely account for the stopping delays. However, in SSTs, when EMG is used to isolate trials where the suppression of movement has occurred (i.e., trials with partial bursts), reaction time delays do not differ significantly between successful ignore and successful stop trials. Instead, the *frequency* of trials in which response delays occurred is greater in stop than ignore trials, accounting for the larger effect in behavioural data observed in prior studies (Weber et al., 2023). Here, we test whether this finding can be generalised to other stopping paradigms (ARI tasks) by including versions of both the SST and ARI task which include ignore trials.

In summary, the current research uses EMG to explore differences in action initiation and cancellation, in motor- and stimulus-selective stopping tasks, with the following goals: **1**) to compare EMG measures of reactive inhibition (CancelTime) between SST and ARI tasks; **2**) to assess whether the re-initiation of unimanual movement following the successful inhibition of bimanual movement is faster during the interruption of anticipated movements (in the ARI task) than reactive movements (in the SST); and **3**) to determine whether previous findings regarding the contribution of attentional processes to stopping delays generalise across task contexts and occur during the inhibition of anticipated movements.

## 2. Methods

### 2.1. Participants

34 healthy adults took part in the study, though the data from 1 participant was excluded due to noise in the EMG data, precluding determination of EMG measures. Thus, data from 33 participants (mean age = 33.3 years, SD = 12.7 years, range = 18-55; 17 male, 16 female) was analysed. Participants were recruited via the University of Tasmania’s School of Psychological Sciences research participation system. As compensation for their time, participants received either two hours of course credit (for those requiring credit) or a $20 shopping voucher. All participants provided informed consent and had normal or corrected to normal vision. This research was approved by the Tasmanian Human Research Ethics Committee.

### 2.2. Computer and response interfaces

Participants were seated approximately 80cm from a computer monitor with forearms and hands resting on a desk shoulder width apart; each index finger rested against one of two custom made response buttons. The buttons were mounted in the vertical plane, such that registering a response required participants to abduct their index fingers (inwards) with the first dorsal interossei (FDI) muscle acting as a primary agonist. This set up maximised FDI activation during voluntary button presses (c.f. finger flexion) and provided high signal-noise ratio in electromyographic data, facilitating the analysis of muscle bursts. Button presses were registered via The Black Box Toolkit USB response pad and recorded using PsychoPy3 (Peirce et al., 2019). The total duration of the experiment was approximately two hours.

### 2.3. Electromyography (EMG)

EMG data were recorded using the computer software Signal (Cambridge Electronic Design Ltd.). EMG recordings were made using disposable adhesive electrodes placed on the first dorsal interossei (FDI) of participants’ left and right index fingers. Two electrodes were placed in a belly-tendon montage on each hand, with a third ground electrode placed on the head of the ulna. The analogue EMG signals were band-pass filtered at 20-1000 Hz, amplified 1000 times, and sampled at 2000Hz. Participants were asked to relax their hands as much as possible between trials, and only activate muscles to press the buttons. Once the experiment commenced, if recordings became noisy with background activity, the researcher would remind the participant to relax their hands.

### 2.4. Behavioural tasks overview

The experiment consisted of four conditions, all of which were developed and run using Psychopy3 (Peirce et al., 2019). Two of these were different variants of the ARI task, developed by modifying a freely available Selective Stopping Toolbox (SeleST; Wadsley et al., 2022). The other two tasks were variants of the stop signal tasks (SSTs). While SeleST does include a stop signal task, this was not appropriate for our current experimental goals, so the SSTs were programmed from scratch. All participants completed the four conditions in a single session. The conditions, explained in detail below, were **1**) stop signal task (SST); **2**) stop signal task with ‘ignore’ trials (SSTignore); **3**) anticipated response inhibition (ARI) task; and **4**) anticipated response inhibition task with ‘ignore’ trials (ARIignore). The order of the four conditions was counterbalanced across participants to minimise practice and fatigue effects. Each condition was divided into 60-trial blocks (with the number of blocks varying depending on condition); rest breaks were provided between blocks and on completion of each condition. During the breaks, text appeared asking participants to rest their hands for a moment and to press one of the buttons when they were ready to continue. Below this text message, feedback was displayed indicating the proportion of correct responses and average reaction time (RT) of correct responses from the previous block. The length of these breaks was left to the participants’ discretion. Trial order within each block was pre-randomised, with the sole criterion that no more than two infrequent trial types (stop or ignore trials; described below) occurred consecutively. The same pre-randomised trial order was used for the ARI and SST to minimise any potential order effects from impacting comparisons between conditions. A different sequence (that included ignore trials) was used for both the SSTignore and the ARIignore conditions. Every trial, in all tasks, were separated by a 1000ms inter-trial interval.

### 2.5. Stop signal tasks (SST and SSTignore)

Prior to completing either of the SSTs participants completed a 20-trial block consisting of only go trials. Go trials began with two white parallel, vertical bars appearing for a pre-randomised duration drawn from a truncated exponential distribution ranging from 250-1000ms. This variable pre-cue period served to reduce the predictability of the subsequent go signal onset and ensure that participants were responding to the visual stimulus, rather than predicting its appearance (Verbruggen et al., 2019). Following this fixation (pre-cue) period, the white bars would turn black, which acted as the go signal and participants were required to enact a bimanual movement, pressing both the left and right buttons, simultaneously and as quickly as possible. To encourage fast responses, feedback was provided at the end of each trial based on how quickly the participant responded, and this was accompanied by a scoring system. The scoring system, and the trial-level coloured feedback, was explained to the participant in instruction slides that appeared prior to the task. During the initial 20-trial block of go trials, RTs of <= 200ms would result in 100 points and green boxes appearing around the stimuli; RTs > 200ms and <= 250ms would result in 50 points and yellow boxes appearing around the stimuli; and RTs > 250ms would result in 20 points and orange boxes appearing around the stimuli. The points were only displayed to the participant at the end of the block and during breaks. Following the initial Go-only block, the participant’s mean RT on successful trials (meanGoRT)would act as a new threshold for the scoring system during the main SSTs, thus providing an individualised RT benchmark. As such, during the SST and SSTignore, RTs of <= meanGoRT would result in 100 points and green boxes appearing around the stimuli, RTs > (meanGoRT+50ms) and < (meanGoRT+100ms) would result in 50 points and yellow boxes appearing around the stimuli, and RTs >= (meanGoRT+100ms) would result in 20 points and orange boxes appearing around the stimuli. If participants made no response, no points were awarded and red boxes would appear around the stimuli. This feedback and scoring system was included to minimise the degree to which participants would slow their responses in anticipation of stop signals (i.e., proactive slowing), a behavioural adjustment which can interfere with the staircase algorithm resulting in very high accuracy rates and invalidating SSRT calculations (Verbruggen et al., 2019).

The SST was comprised of 180 trials (120 go trials, 30 stop-left trials, and 30 stop-right trials) across 3 blocks. During stop trials, after the presentation of the go signal and following a variable “stop signal delay” (SSD; explained below), one of the bars would change colours to either cyan or magenta. The colour used for stop signals was counterbalanced between participants to mitigate any confounding colour effects. However, the stop signal colour did not vary between conditions for a single participant. This colour change acted as the stop signal and indicated that the participant should attempt to cancel their response on the side where the colour change has occurred (while continuing their response with the other hand). A trial was considered correct when participants pressed *only* the button on the side on which the stop signal *did not* occur (that is, they successfully inhibited the response on the side on which the stop signal occurred). Feedback on the side of the stop signal would be green for correctly inhibited responses, or red for incorrect responses, while feedback on the non-stopping side operated by the same rules as described above for go trials. As well as the coloured feedback, text would appear reading “Correct” for correct stop trials and “Incorrect” for any other response (including if the participant refrained from pressing both buttons or inhibited the *wrong* response). This additional text feedback was included to minimise the potential confusion of coloured feedback alone.

SSD refers to the time between the presentation of the go signal and the stop signal. Separate SSDs were used for stop-left and stop-right trials. Initially, both SSDs were set to 200ms, and this was adjusted using a staircase algorithm based on success in the previous trial of that condition. For example, following a correct stop-right trial, the SSD for the subsequent stop-right would increase by 50ms, making stopping more difficult. Following a failed stop-right trial, the SSD would decrease by 50ms, making a subsequent stop-right trial easier. Using this staircase procedure, the probability of successfully stopping approximates to 50%, enabling assessment of both successful and failed stop trials, as well as calculation of SSRT (Verbruggen et al., 2019). The minimum and maximum SSDs were 50ms and 400ms, respectively.

The SSTignore condition was comprised of 360 trials (240 go trials, 30 stop-left trials, and 30 stop-right trials, 30 ignore-left trials and 30 ignore-right trials; 6 blocks). This task followed the same procedure as the SST, with the addition of ‘ignore’ trials. As with the stop trials, ignore trials involved one of the bars changing colour following the go signal. For half of the participants the colour assigned to ignore signals was magenta and for half it was cyan (whichever colour was *not* assigned to the stop trials for that participant was used for the ignore trials). During these trials, participants were required to ignore the colour change and continue with the button press (i.e., respond bimanually as though both bars had remained black). The timing of the ignore signals was determined from the SSD value that was set following the previous stop trial occurring on the same side (i.e., the timing of an ignore signal for an ignore-left trial was the same as the SSD set following the last stop-left trial). Performance on ignore trials did not influence SSDs. The proportion of trials involving unexpected signals (stop *or* ignore signals) remained consistent with that in the SST (where 1/3 of trials were stop trials). i.e., in this variant of the task, 1/6 of all trials were stop trials and 1/6 of all trials were ignore trials

### 2.6. Anticipated response inhibition tasks (ARI and ARIignore)

Prior to completing either of the ARI tasks participants completed a 20-trial block consisting of only go trials. At the beginning of each go trial two vertically aligned white rectangles appeared on the screen which were intersected by a horizontal black “target” line appearing two-thirds of the way up the rectangles (refer to Figure 1). After 750ms the white rectangles would begin to fill from the base, gradually changing colour to black. This rising colour change moved at the same rate for both the left and right rectangles, intersecting the target line 800ms after filling began, and reaching the top of the rectangle at 1200ms. The left and right rectangles were related to the participants’ left and right hands, respectively. Participants were required to coincide their presses with the left and right buttons to the moment when the rising colour change intersected the horizontal target line (800ms from when the trial began). When a button was pressed, the rising colour change on the side of the button press would stop rising and remain stationary for the rest of that trial. Following the 1200ms response window, feedback was provided by way of the target line changing colour, based on how close the bar was to the target line when the participant pressed the button. As with the SSTs, this was accompanied by a scoring system. If the response was made within 25ms of the target line (i.e., more than 25ms before the target, less than 25ms after the target) it would change colours to green and 100 points were awarded. Responses between 25-50ms of the target line would result in the bar turning yellow and 50 points being awarded. Responses beyond 50ms of the target line would result in the bar turning orange and 25 points being awarded. As with the SSTs, this scoring system was conveyed to the participant in instruction slides appearing prior to the task beginning and the participants’ score was displayed during breaks and at the end of each experimental block.

**Figure 1:**
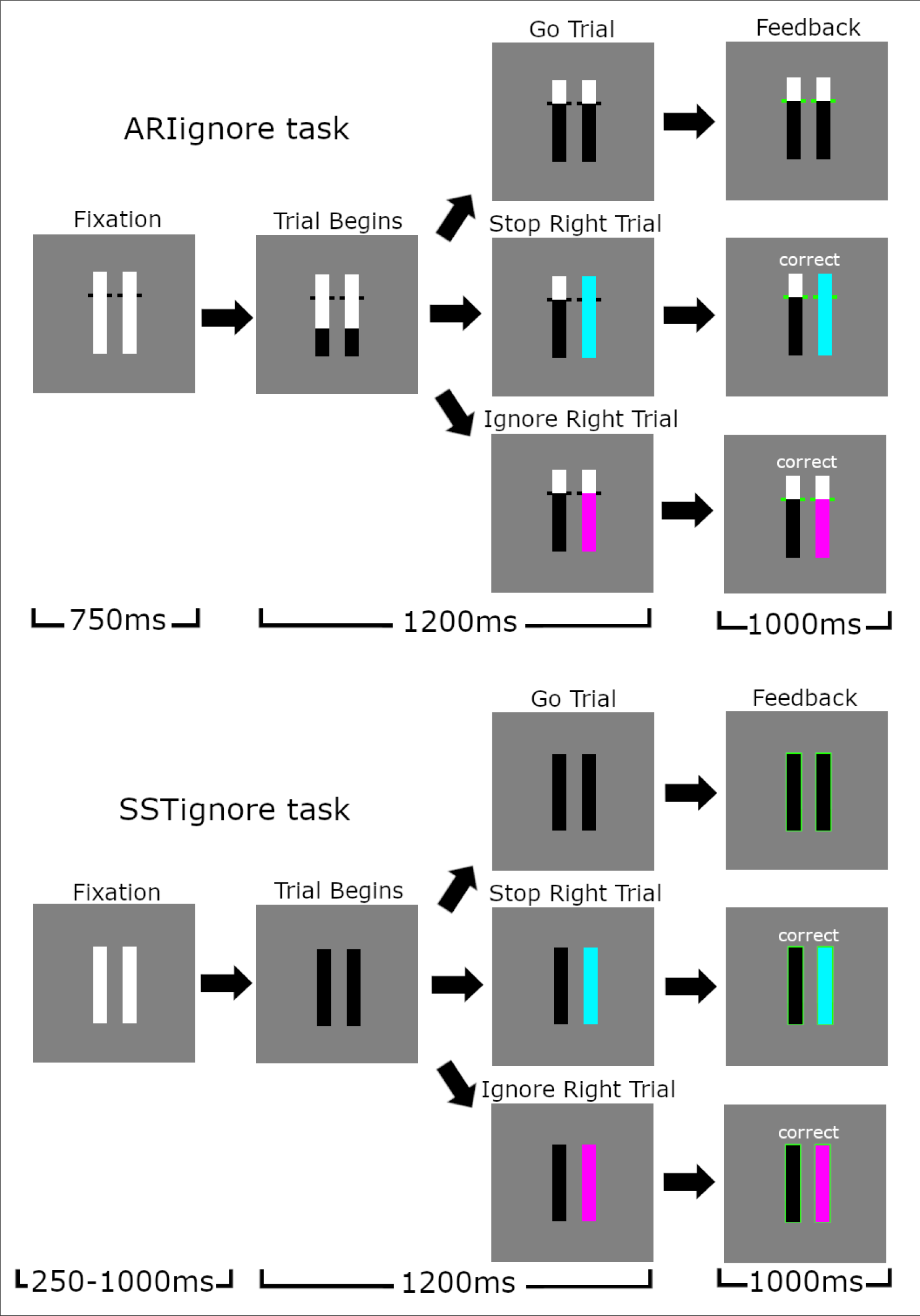
Trial types in the SSTignore and ARIignore conditions*. The bars appearing on the left and right of the screen were associated with responses with the left and right hands, respectively. Stop and ignore signals could occur on either the left or right side (with equal likelihood), although for demonstration purposes only stop right and ignore right trials are shown here. Feedback for correct responses is shown (see text for feedback on incorrect responses). The colours for stop and ignore stimuli are depicted as cyan for stop and magenta for ignore, although this was counterbalanced across participants (see methods). * NB: The trial sequences in the SST and ARI conditions were identical as those depicted here but without ignore trials.

The ARI task was comprised of 180 trials (120 go trials, 30 stop-left trials, and 30 stop-right trials; 3 blocks of 60 trials) and used the same trial order as the SST. During stop trials, one of the rising black stimuli bars would change colour to either cyan or magenta (depending on counterbalancing group) *prior* to reaching the target line. This colour change acted as the stop signal, indicating that the participant should attempt to cancel their response on the side where the colour change has occurred (while continuing their response with the other hand to try and stop the bar accurately at the target line). As with the SSTs, a stop trial was considered correct when participants pressed *only* the button on the side which the stop signal *did not* occur. On the stopping side, the colour of the feedback (conveyed by the target bar changing colour) would be green for correctly inhibited responses, or red for incorrect responses. Additionally, text would appear reading “Correct” for correct responses and “Incorrect” for any other response. In the ARI tasks we refer to the time between the stop signal occurring and when the rising bar intersects the target (at 800ms) as the SSD. As with the SSTs, the SSD in the ARI tasks used a separate tracking procedure for left and right hands. The SSD for both hands began at 300ms (i.e., 500ms from the beginning of the trial, when the bars started rising) and this was adjusted based on success in the previous stop trial: Following a correct stop-right trial the SSD for the subsequent stop-right would decrease by 25ms, making it harder to stop on the next stop trial. Following a failed stop-right trial the SSD would increase by 25ms, making it easier to stop on the next trial. Smaller steps were used in the ARI tasks than the SSTs (which used 50ms adjustments) due to the fact that ARI tasks demonstrate less variability in RT distributions (Leunissen et al., 2017), allowing for a more fine-grained approach.

The ARIignore condition was comprised of 360 trials (240 go trials, 30 stop-left trials, and 30 stop-right trials, 30 ignore-left trials and 30 ignore-right trials; 6 blocks of 60 trials) and used the same pre-randomised trial order as the SSTignore. This task followed the same procedure as the ARI task, with the addition of ‘ignore’ trials. As with the stop trials, ignore trials involved one of the bars changing colour following the go signal (to either cyan or magenta; whichever colour was not used for stop trials for that participant). During these trials, participants were required to ignore the colour change and continue with the button press (i.e., respond bimanually when the rising bars reached the target). The timing of the ignore signals was determined from the SSD value that was set following the previous stop trial occurring on the same side. Performance on ignore trials did not influence SSDs.

### 2.7 Data Analysis – Behavioural

Statistical analyses were conducted using linear mixed models (LMMs) and generalised linear mixed models (GLMMs) run within the statistical package Jamovi (The Jamovi Project, 2021) which operates via the statistical language R (R Core Team, 2021). GLMMs were run using the Jamovi module GAMLj (Gallucci, 2019). For each model a maximal random effects structure was initially used. If this failed to converge we used a stepwise approach to model simplification whereby we first removed interaction terms from the random effects structure, followed by slope terms (Barr et al., 2013).

#### 2.7.1 Proactive Slowing

When anticipating the possible need to inhibit an upcoming movement, adjustments may be made within inhibitory networks to facilitate the potential upcoming action cancellation (Lavallee et al., 2014). These changes are referred to as proactive inhibition. A behavioural manifestation of these changes is proactive slowing, which is quantified by comparing a participant’s reaction time (RT) in go trials during a task involving stop trials to RTs to go signals during a block consisting only of go signals (van de Laar et al., 2011). Here, we sought to assess proactive inhibition in each of the four stopping tasks. Prior to analysis, we observed that the reaction time distributions in ARI tasks and SSTs precluded the direct comparison of proactive slowing within the one model-based analysis (the distribution of go trial RTs from each condition can be seen in Figure 2). RTs commonly possess a positive skew, which can be accounted for by using a gamma distribution with a log link function (Lo & Andrews, 2015). While this approach is appropriate for the RT data from the SSTs, the distribution on the ARI tasks lacks the positive skew characteristic of gamma distributions, and we thus used a more traditional Gaussian distribution (identity link). As such, two models were run, one comparing go trial RTs in the ARI tasks, another comparing go trial RTs in the SSTs. The final (converging) model for the ARI tasks included the fixed factor of condition (Go Only ARI, ARI, and ARIignore), and the covariate of age. The random effects structure included participant intercepts and random slopes for condition. Similarly, the final converging model for the SSTs included the fixed factor of condition (Go Only SST, SST, and SSTignore), and the covariate of age. The random effects structure included participant intercepts and random slopes for condition.

**Figure 2:**
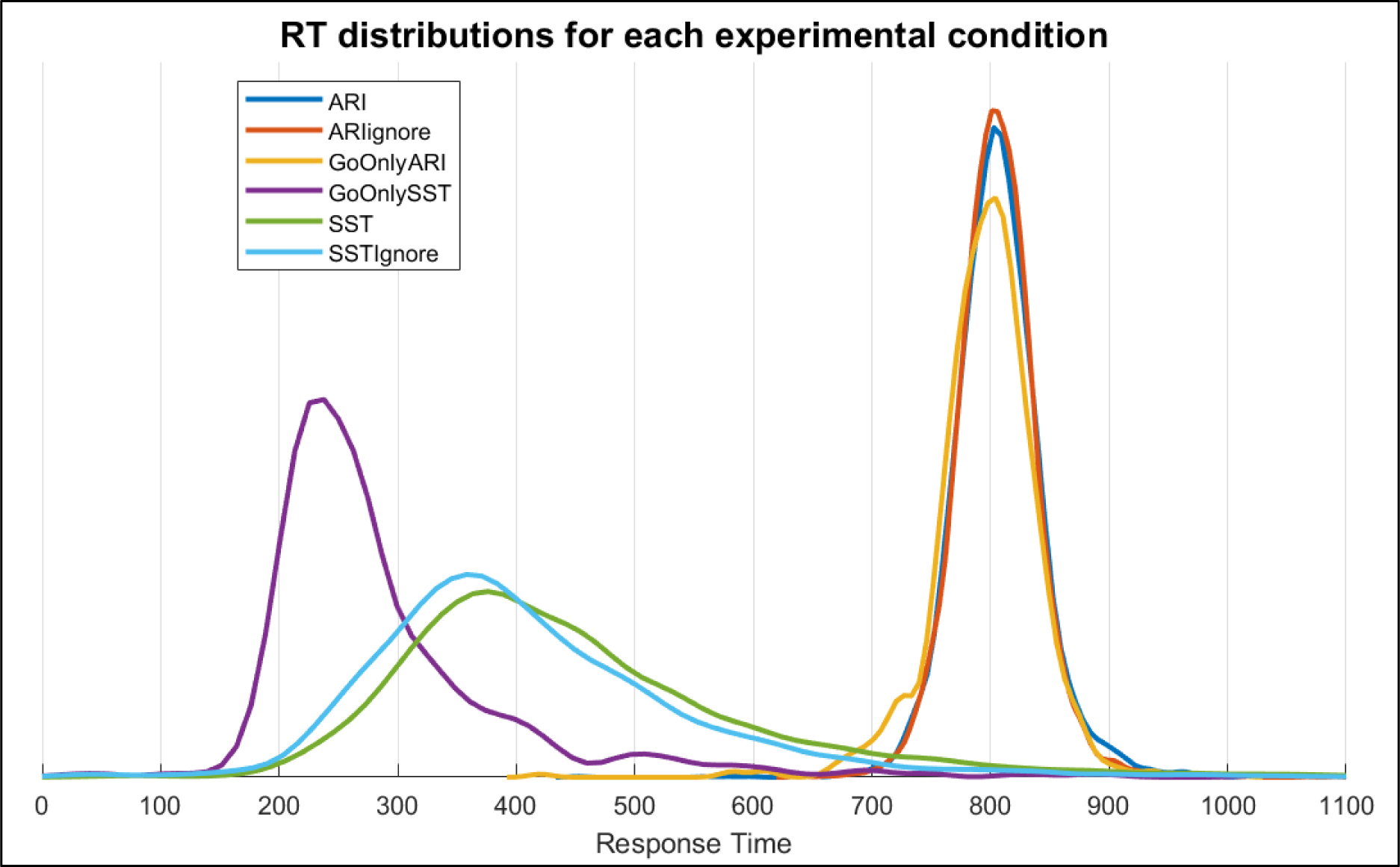
Density plot of Go trial RTs in all six conditions. Note that the y-axis label is omitted here as the trial densities are relative to each condition. This was done to allow for easier visual comparison between the distributions of GoOnly conditions (GoOnlyARI and GoOnlySST; in which fewer go trials were completed) to other conditions. RTs in the ARI tasks centre around 800ms as this was the time point at which the stimuli indicated that a response should be made (refer to section 2.6).

#### 2.7.2 Selective stopping reaction times (stopping delays)

Previous assessments of RT differences in go and successful selective stop trials have involved comparisons of pooled (from all participants) reaction time data recorded from the non-signalled hand in stop trials to mean reaction times in go trials. Here, we sought to compare stopping delays between tasks. As such, we chose to determine a trial-level measure which could be compared across task contexts. To do this, each participant’s mean Go trial RT was determined in each condition. This was then subtracted from RTs from the non-signalled hand in successful stop and ignore trials^1^ within that condition. The resulting data was analysed using an LMM (Gaussian distribution and identity link function). The model included condition (SST, SSTignore, ARI, ARIignore) and trial type (stop, ignore) as fixed factors, and the covariates of age and SSD. In ARI tasks, longer SSDs occur earlier in the trial and are easier to stop to, whereas in SSTs longer SSTs occur later in the trial and are harder to stop to. To compensate for this difference SSDs in the ARI tasks were inverted by subtracting the SSD from the target response time (800ms). The random effects structure included participant intercepts and slopes for both condition and trial type.

#### 2.7.3 SSRT

SSRT was calculated using the integration method (Verbruggen et al., 2019). The horserace model upon which this method is based assumes that mean RTs on unsuccessful stops are faster than go trial RTs. In the ARI and ARIignore tasks, no single participant met this requirement in both of the tasks (5 participants met this requirement for the ARI task, and another 2 participants met this requirement for the ARIignore). Similarly high levels of exclusion have been noted in previous motor-selective ARI tasks (e.g., Wadsley et al., 2022, where only 20% of participants met this assumption). As such, here we only include SSRT results from the SSTs (for EMG based calculations of inhibition latency in both ARIs and SSTs refer to section 3.5). Three participants were excluded due to not meeting this requirement in the SSTignore condition, but no exclusions were required for the SST. Unlike our other analyses which utilise trial-level data, each participant has a single SSRT score for each condition. As such, a general linear model was run, with SSRT as the dependent variable, condition (SST, SSTignore) as an independent variable, and age as a covariate.

### 2.8 Data Analysis – EMG

#### 2.8.1. Processing of EMG data

EMG data processing was performed using MATLAB (MathWorks, 2018). EMG signals were digitally filtered using a fourth-order band-pass Butterworth filter at 20–500 Hz. The precise onset and offset times of EMG bursts were detected using a single-threshold algorithm (Hodges & Bui, 1996): Signals from each trial were rectified and low-pass filtered at 50 Hz. Following this, we used a sliding window of 500ms to find the trial segment with the lowest root mean squared (RMS) amplitude, which was used as a baseline. EMG bursts were identified when the amplitude of the smoothed EMG was more than 3 SD above baseline. EMG bursts separated by less than 20ms were merged together to represent a single burst.

Following identification of the onset and offset times, we used time constraints to identify two types of EMG burst: The *RT-generating burst* was identified as the last burst with an onset occurring *after* the go signal and *prior* to the recorded button press. Some trials also exhibited *partial bursts*, i.e., EMG bursts that started to decrease in amplitude before enough force was generated to trigger an overt button press. Partial bursts were identified in each hand as the earliest burst where peak EMG occurred after the presentation of the stop/ignore signal but prior to the onset of the RT-generating burst from the responding hand. Furthermore, peak EMG amplitude was required to be greater than 10% of the average peak amplitude from that participant’s successful go trials in that condition. Critically, these time and amplitude constraints exclude activity occurring *after* the RT-generating burst, as this is likely related to mirror activity, or other activity unrelated to the task. For each burst (RT-generating and partial), we extracted the time of burst onset and offset, as well as the time of peak amplitude.

EMG envelopes were obtained using full-wave rectification and a low-pass filter at 10 Hz. EMG envelopes were used to extract the peak amplitude. For each subject, the amplitude of EMG envelopes from each hand was normalised by the average peak EMG from successful bimanual go trials in the SST condition, thus allowing direct comparisons between participants and conditions.

#### 2.8.2. Frequency of partial bursts

We can only assesses CancelTime in the subset of successful stop trials *with* partial responses, and thus the measure is “censored” at both ends of the distribution: Inhibition occurring “too late” is censored as we exclude failed stop trials, whereas early inhibition results in successful stop trials without partial bursts. As such, the calculation of inhibition latency via EMG depends on there being sufficient trials with partial responses to justify the use of CancelTime as a reliable measure, and a larger proportion of trials with partial responses reduces the censoring effects. For an in-depth description and modelling of censoring effects, see Salomoni et al. (2023). Here, we sought to determine whether the selective ARI or selective SSTs were more or less conducive to producing partial responses. A GLMM with a binomial distribution and probit link function was run using a categorical dependent variable reflecting the presence/absence of partial burst in either hand. The independent variables of experiment (SST, SSTignore, ARI, ARIignore) and trial type (stop, ignore) were used, and age was included as a covariate. The final model featured a random effects structure of participant intercepts and slopes for experiment and trial name.

#### 2.8.3. CancelTime

On successful stop trials where a partial burst is observed, the latency of the peak amplitude of the partial EMG burst relative to that trial’s SSD can be used to determine a trial-level estimate of stopping latency (Jana et al., 2020; Raud et al., 2022). Past research has termed this CancelTime (Jana et al., 2020). CancelTime is conventionally calculated based on partial bursts in the stopping hand. Here we expand upon this definition, calculating CancelTime based on partial bursts in either hand, (or where they have been detected in both hands, whichever partial burst began to decrease in amplitude at an earlier time point). This more flexible approach is made possible due to the use of a selective stopping task: either or both hands may demonstrate movement suppression, whereas previous papers have calculated CancelTime in simple stopping tasks where only the one movement is being made, or based solely on the stopping hand (Jana et al., 2020; Raud et al., 2022). Furthermore, our algorithm for detecting partial bursts prevents the spurious inclusion of RT generating activity in the non-cued (responding) hand by merging partial bursts which are separated by less than 20ms together (i.e., a complete suppression of muscle activity followed by a reactivation must have occurred for the partial burst to be considered distinct from the RT generating burst). Accordingly, we can accurately detect partial bursts in both the stopping and continuing hands. Indeed, our recent model of selective stopping (Gronau et al. 2023) is consistent with a global (bilateral) inhibition occurring prior to initiation of the non-cancelled, selected response, such that bilateral partial bursts may occur. Because the response window following SSD could vary between tasks, an upper cut-off of 2SD above the mean CancelTime was used to remove potential outliers (70 trials removed; 5.29% of trials). A GLMM with a gamma distribution and a log link function was run on this trial-level data, using experiment (SST, SSTignore, ARI, ARIignore) as an independent variable and age as a covariate. The random effects structure included participant intercepts and slopes for experiment.

#### 2.8.4. Selective stop RTs in the presence/absence of partial bursts

Our previous research investigating inhibitory responses to stop and ignore signals found no significant differences in RTs between successful selective stop trials and ignore trials when we compared trials in which partial bursts were observed (Weber et al., 2023). Here, we sought to investigate whether this finding generalised to the inhibition of anticipated responses. As such, an additional model was run using trial-level stopping delays (see 2.7.2) as the dependent variable. The independent variables were experiment (SSTignore, ARIignore), trial type (stop, ignore) and partial burst (present/absent), with age included as a covariate. The random effects structure included participant intercepts and slopes for each of the three independent variables.

#### 2.8.5. Speed of action reprogramming

Following visual inspection of the EMG profiles from each task, we conducted an analysis to determine whether the speed of unilateral action initiation following suppression of a bimanual movement in correct stop and ignore trials with partial bursts differed between experimental conditions (i.e., whether the time between the suppression of the movement in the non-signalled hand and the initiation of the unimanual movement differed between trial types and tasks). We used the duration of time between the peak of the partial burst (CancelTime) and the initiation of the RT generating burst (RT-generating burst onset time) as a trial-wise dependent variable. We imposed an upper cut-off of 400ms (removing 1.69% of trials) to ensure that the longer response window (following SSD) in the SSTs (c.f. ARI tasks) didn’t influence results. This analysis was run using data from the non-signalled hand in each condition (e.g., left hand in a right-stop trial). The data were analysed using a GLMM with a gamma distribution with a log link function. Condition (SSTignore, ARIignore) and trial type (stop, ignore) were included as independent variables and age was included as a covariate. Random effects included participant intercepts and random slopes for condition and trial type.

## 3. Results

### 3.1. Behavioural Results

#### 3.1.1. Accuracy

Trial success rates indicated that participants completed the task well, with few errors on go and ignore trials; moreover, stopping success rates between 46-55% for all tasks indicate that the SSD staircasing procedure was effective (percentages of correct trails for each trial type in each condition are presented in Table 1).

**Table 1:**
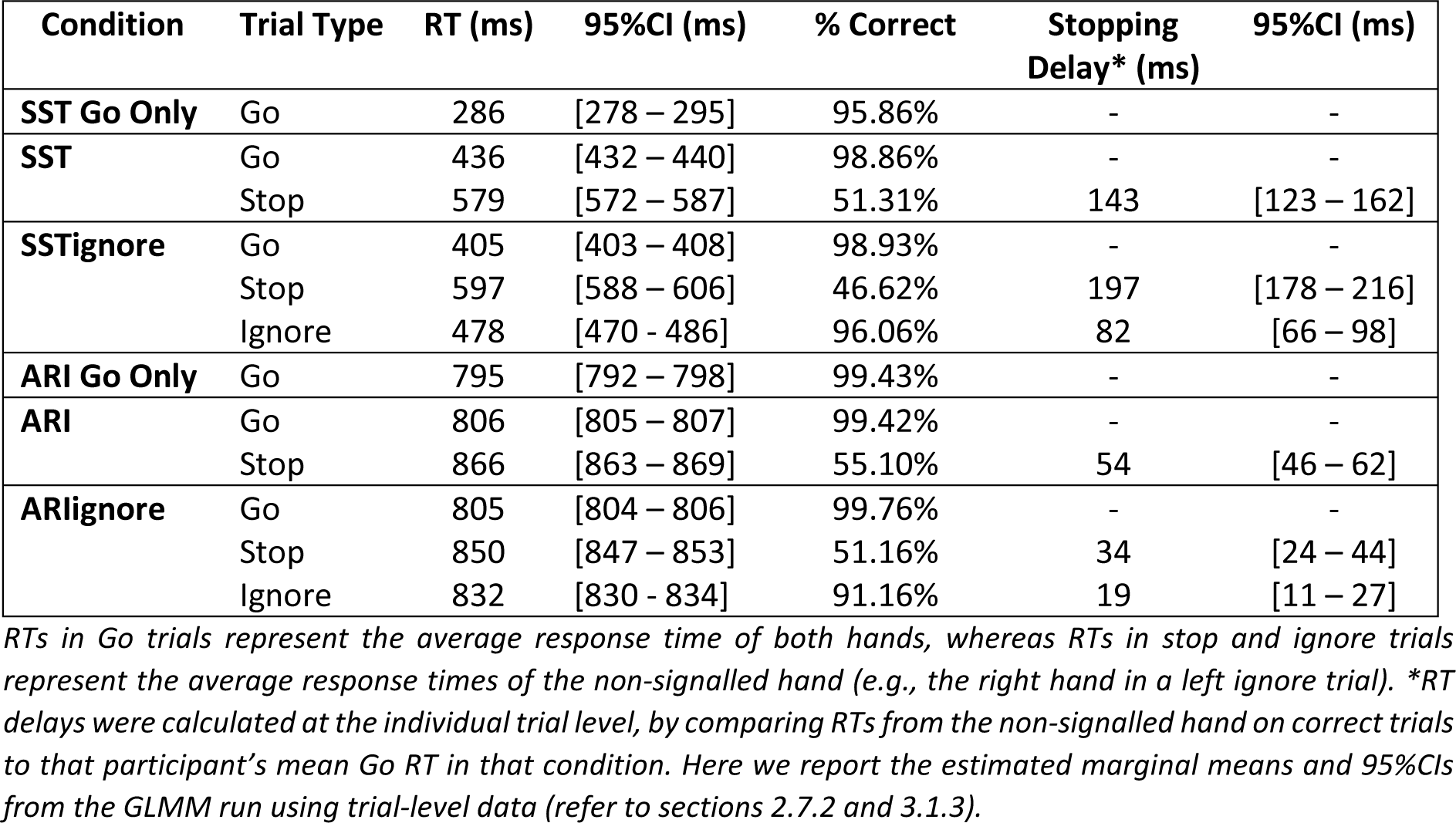
Key behavioural results.

#### 3.1.2. Proactive Slowing

Mean Go trial RTs are shown in Table 1, whereas RT distributions from Go trials across all six conditions are depicted in Figure 2. The proactive slowing analysis for the SSTs revealed a statistically significant main effect of condition ꭓ^2^(2) = 121.34, *p* < 0.001. The covariate of age was not statistically significant ꭓ^2^(1) = 3.05, *p* = 0.08. Bonferroni corrected post-hoc tests revealed significant proactive slowing effects (i.e., slowing relative to Go Only condition) in the SST (*z* = 10.84, *p* <0.001, *d* = 1.23) and the SSTignore (*z* = 10.45, *p* <0.001, *d* = 1.01). The post-hoc tests also revealed RTs in the SST were significantly slower than those in the SSTignore (*z* = 3.77, *p* <0.001, *d* = 0.23).

The comparable analysis for ARI conditions also revealed a significant main effect of experimental condition *F*(2,32) = 9.31, *p* < 0.001. The covariate of age was not statistically significant *F*(2,31) = 0.79, *p* = 0.38. Bonferroni corrected post-hoc tests revealed small but significant proactive slowing effects in the ARI (*t* = 4.21, *p* <0.001, *d* = 0.29) and the ARIignore (*t* = 4.16, *p* <0.001, *d* = 0.27) relative to the GoOnly ARI. No significant difference in mean RT was observed between the ARI and ARIignore tasks (*t* = 0.80, *p* = 1.000, *d* = 0.03). Visual inspection of the RT distributions suggests that the significant effect was driven by some early responses in the Go Only ARI condition, rather than proactive slowing in the conditions with stop signals. Given that participants completed the go only conditions prior to the main blocks (as they were used to determine baseline RTs), this result may have been influenced by practice effects.

#### 3.1.3. Selective stopping reaction times (stopping delays)

The GLMM revealed significant main effects of condition *F*(3,41) = 54.53, *p* < 0.001, trial type *F*(1,33) = 277.96, *p* < 0.001, and a significant interaction between condition and trial type *F*(1,7639) = 324.43, *p* < 0.001. The covariate of age was not statistically significant *F*(1,31) = 1.26, *p* = 0.271, β = 0.27 (95%CIs -0.19, 0.73). The covariate of SSD revealed that longer SSDs were associated with larger stopping delays *F*(1,4995) = 58.05, *p* < 0.001, β

= 0.16 (95%CIs 0.12, 0.20). Bonferroni corrected post-hoc tests revealed stopping delays in ignore trials were significantly smaller than those in stop trials in the SSTignore, *t*(75) = 23.81, *p* < 0.001, *d* = 0.80. The same comparison was also significant in the ARIignore *t*(71) = 3.25, *p* = 0.049, *d* = 0.34. Stopping delays in ignore trials were larger in the SSTignore than in the ARIignore *t*(40) = 6.95, *p* < 0.001, *d* =0.51. Stopping delays in stop trials were larger in the SSTignore than in SST (*t*(38) = 8.14, *p* < 0.001, *d* = 0.51), but stopping delays in stop trials in the ARIignore task were smaller than those in the ARI task (*t*(149) = 4.41, *p* < 0.001, *d* = 0.42). Estimated marginal means are presented in Table 1.

#### 3.1.4. SSRT

The GLM on SSRT (conducted only on SST and SSTignore conditions – see methods) revealed the main effect of condition *F* (1,57) = 1.08, *p* = 0.303 was not statistically significant; Mean SSRT in the SST condition was 214ms, 95%CIs [198 – 230ms] while mean SSRT in the SSTignore condition was 225ms, 95%CIs [210 – 241ms].The covariate of age was significant *F*(1,57) = 5.28, *p* = 0.025, η^2^ = 0.08, with greater age being associated with a higher SSRT, β = 1.01 (95%CIs 0.13, 1.88).

### 3.2. EMG results

#### 3.2.1. Frequency of partial bursts

The percentages of successful stop and ignore trials with partial bursts (split by trial type and condition) are presented in Table 2. The GLMM examining percentage of trials with partial bursts revealed a significant main effect of condition ꭓ^2^(3) = 27.68, *p* < 0.001, a significant main effect of trial type ꭓ^2^(1) = 8.52, *p* = 0.004, and a significant interaction between condition and trial type ꭓ^2^(1) = 36.93, *p* < 0.001. The covariate of age was not statistically significant ꭓ^2^(1) = 1.31, *p* = 0.253. Bonferroni corrected post-hoc comparisons to explore the interaction revealed that this was driven by the SST conditions having a larger proportion of partial bursts in stop trials that the ARI conditions (ARI vs SST, *z* = 3.33, *p* = 0.02; ARIignore vs SST, *z* = 4.57, *p* < 0.01; ARI vs SSTignore, *z* = 4.44, *p* < 0.01), but not in ignore trials (SSTignore vs ARIignore, *z* = 1.20, *p* = 1.00). No significant differences were found between proportions of partial bursts in stop trials in the SST compared to the SSTignore (*z* = 2.64, *p* = 0.23), nor the ARI compared to the ARIignore (*z* = 1.32, *p* = 1.00).

**Table 2:** Key physiological results.

| Condition | TrialType | % Correct with partial bursts | CancelTime (ms) | 95%CI (ms) | Action reprogramming (ms) | 95%CI (ms) |
| --- | --- | --- | --- | --- | --- | --- |
| ARI | Stop | 29.69% | 145.42 | [134.84 – 156.83] | 99.73* | [94.64 – 104.82] |
| ARIignore | Stop | 26.07% | 142.11 | [129.75 – 155.66] | 94.06 | [86.10 – 102.76] |
|  | Ignore | 24.54% | - | - | 110.39 | [101.70 – 119.82] |
| SST | Stop | 39.64% | 136.74 | [126.80 – 147.46] | 132.96* | [126.20 – 139.71] |
| SSTignore | Stop | 43.42% | 144.49 | [136.76 – 152.65] | 137.62 | [127.69 – 148.32] |
|  | Ignore | 29.24% | - | - | 145.46 | [134.65 – 157.13] |

\*Action reprogramming values for the ARIignore and SSTignore conditions represent estimated marginal means from a GLMM run on this data (refer to section 2.8.5). However, given the ARI and SST were not included in this model, action reprogramming values for these conditions represent *descriptive* means. Columns representing percentages of partial bursts and CancelTime are estimated marginal means from associated analyses.

#### 3.2.2. CancelTime

Mean CancelTime values for each condition are presented in Table 2. The GLMM run on CancelTime revealed no significant differences between conditions ꭓ^2^(3) = 2.40, *p* = 0.493 though the covariate of age was statistically significant ꭓ^2^(1) = 4.58, *p* = 0.03, with CancelTime increasing with greater age β = 1.004 (95%CIs 1.000, 1.007).

#### 3.2.3. EMG profiles

Figures 3 and 4 show average EMG profiles for the ARIignore task and the SSTignore task, respectively. Profiles are synchronized to the onset of the RT-generating burst, allowing for clear observation of the partial activations in stop and ignore trials (blue line). Moving the synchronization reference away from the peak of the RT-generating burst causes the combined EMG profiles to “blur”, due to the averaging of profiles whose peaks are not perfectly aligned in time. As a result, the normalised peak amplitude of the average EMG profiles in these figures appears to be below 1. Note that the RT generating burst in partial burst trials (blue) reaches a greater amplitude than responses without a partial burst (green). This indicates greater drive after a partial burst and successful stopping, to overcome the inhibition and respond rapidly unilaterally.

**Figure 3.**
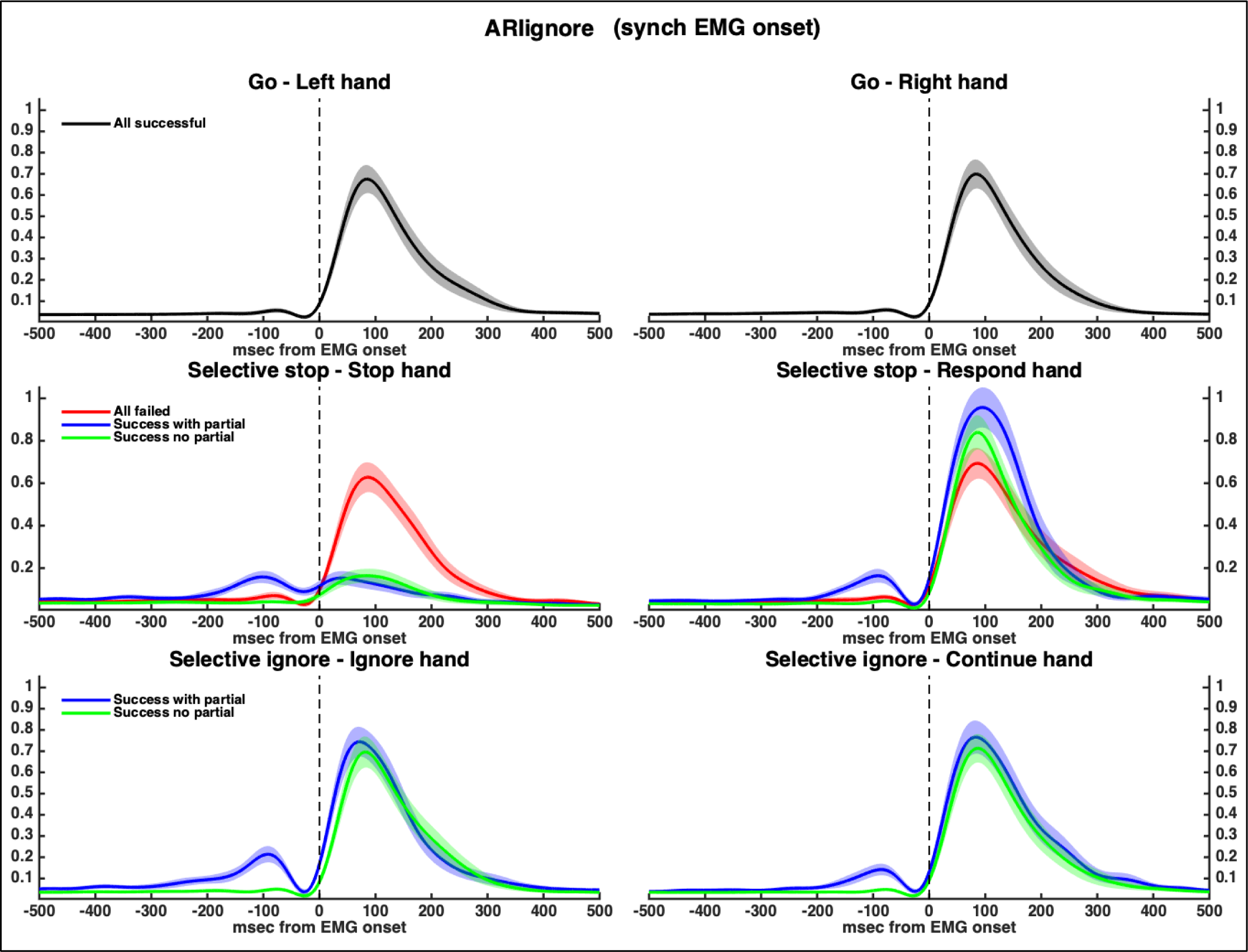
EMG profiles for the Anticipated Response Inhibition task with ignore trials (ARIignore) split trial type, and presence/absence of partial burst. Trials are synchronised to the onset of the RT generating burst in order to allow for observation of the presence of partial activations (and the subsequent inhibition). Shaded areas represent 95%CIs.

**Figure 4.**
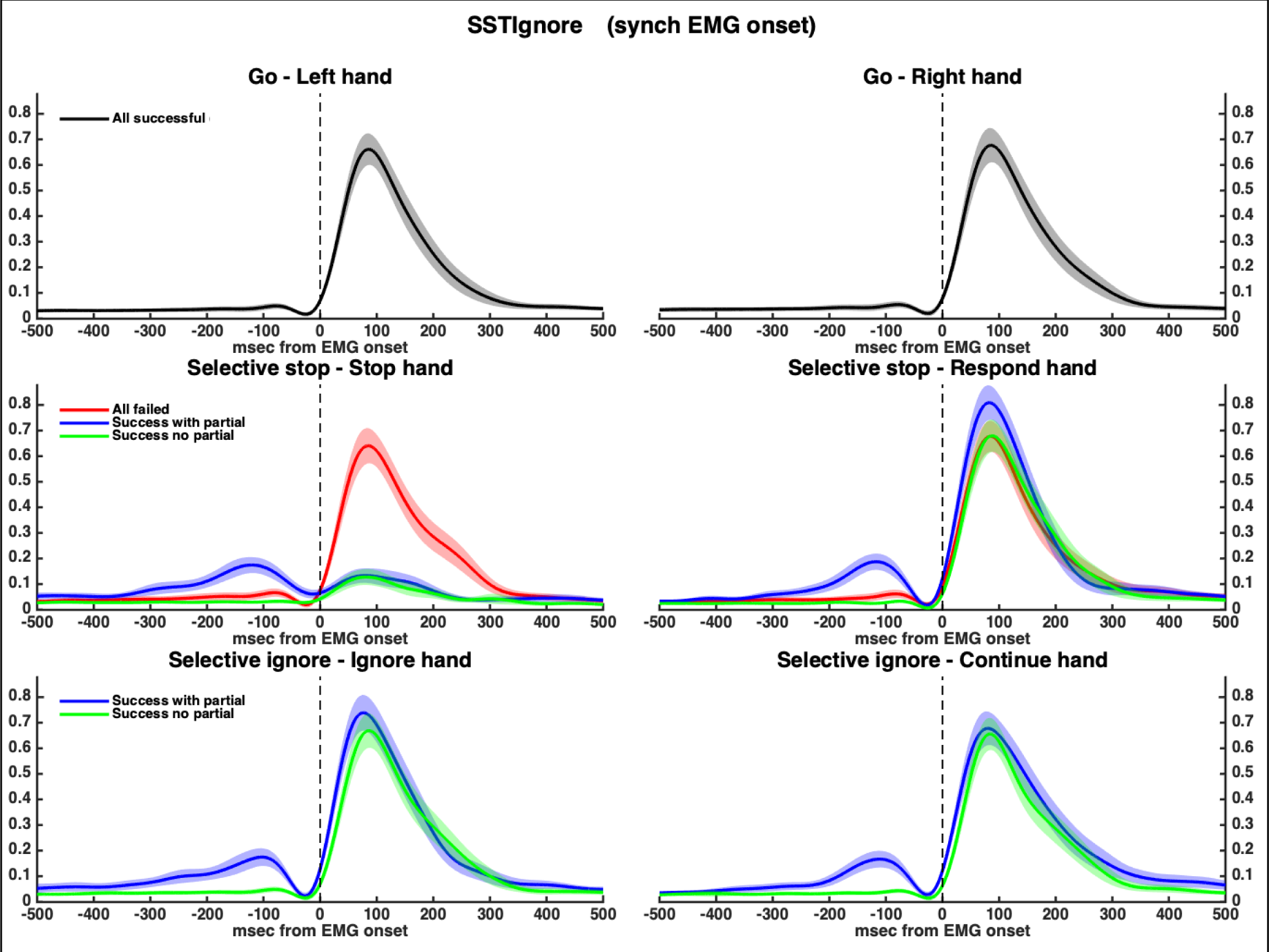
EMG profiles for the Stop Signal Task with ignore trials (SSTignore) split trial type, and presence/absence of partial burst. Trials are synchronised to the onset of the RT generating burst in order to allow for observation of the presence of partial activations (and the subsequent inhibition). Shaded areas represent 95%CIs.

#### 3.2.4. Selective stop RTs in the presence/absence of partial bursts

The results of the model analysing stopping delays split by the presence or absence of partial bursts are depicted in Figure 5. The model revealed significant main effects of condition *F*(1,34) = 164.73, *p* < 0.001, trial type *F*(1,40) = 132.64, *p* < 0.001, and presence/absence of partial burst *F*(1,31) = 147.33, *p* < 0.001. All two way interactions were significant (*p* < 0.001), as was the higher-order three-way interaction *F*(1,5206) = 123.94, *p* < 0.001. The covariate of age was not statistically significant *F*(1,31) = 0.56, *p* = 0.458. Tests of simple main effects were conducted to investigate the three-way interaction, specifically comparing stop and ignore trials with and without partial bursts in both conditions. In the ARIignore condition no significant differences between stop and ignore trials were observed in trials where partial bursts occurred, *t*(572) = 1.02, *p* = 0.306, *d* = 0.17, though in trials without partial bursts significant differences in stopping delays were observed *t*(101) = 3.93, *p* < 0.001, *d* = 0.46. Similarly, in the SSTignore condition no significant differences between stop and ignore trials were observed in trials where partial bursts occurred, *t*(331) = 0.64, *p* = 0.523, *d* = 0.04. though in trials without partial bursts significant differences were observed *t*(151) = 26.92, *p* < 0.001, *d* = 1.10.

**Figure 5.**
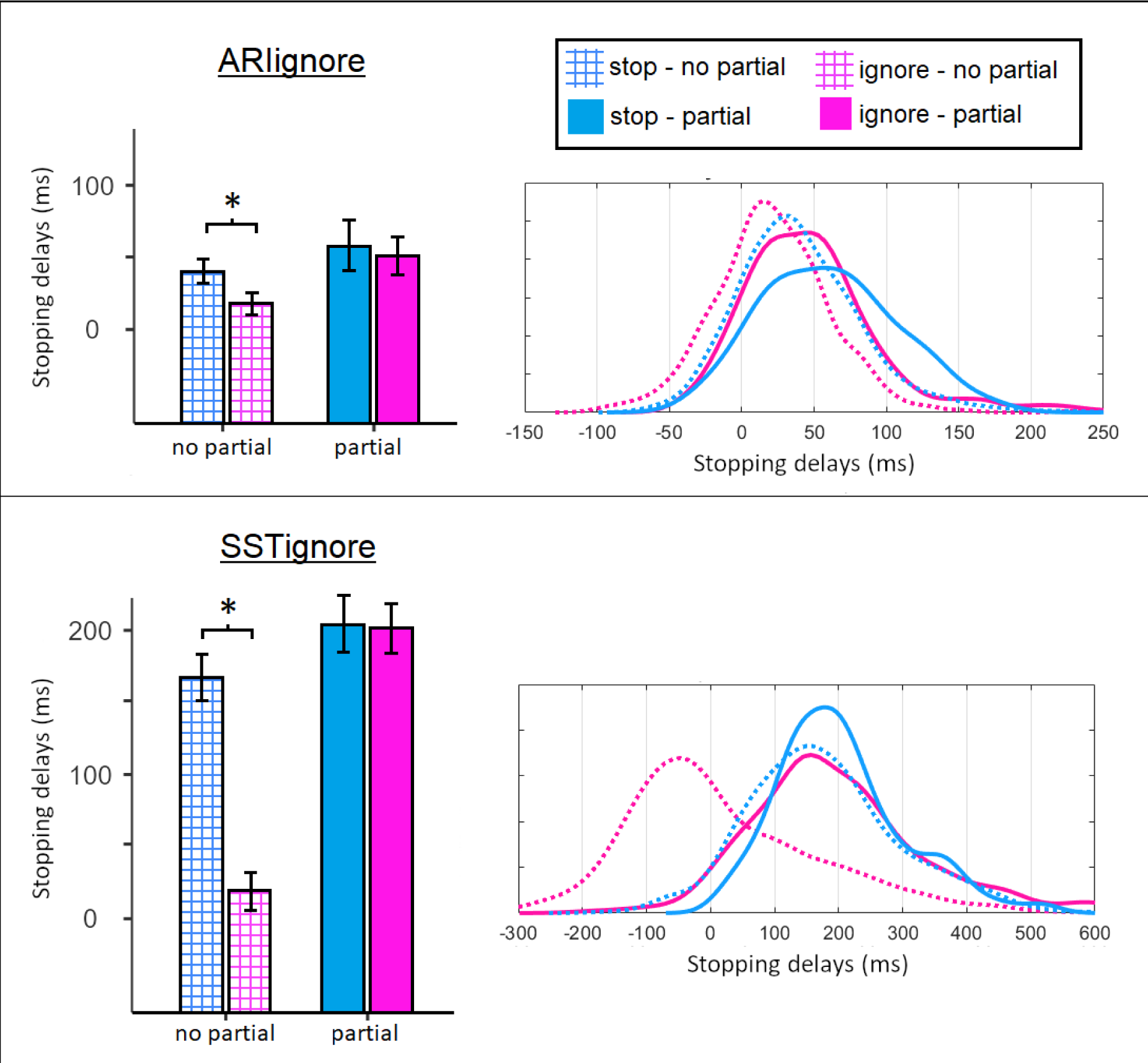
Stopping delays to stop and ignore trials in the SSTignore and ARIignore tasks split by presence/absence of partial burst. * = *p* < 0.001. Error bars in box plots represent 95%CIs. We note that when trials in which partial bursts are detected are compared, no differences in stopping delays can be seen between stop and ignore trials. Figures on the right are density plots representing the distribution of stopping delays. Note that the y-axis label is omitted here as the trial densities are relative to each category, to allow for clearer visual comparisons. In the SSTignore in particular, splitting trials into those with and without partial bursts demonstrates how correct ignore trials conflate fast responses (with no partial burst, where the participant is most likely responding prior to noticing the ignore signals) and slower responses (in which the participant has paused in response to the ignore cue). In stop trials, fast responses are excluded from calculations of stopping delays, as these are counted as incorrect.

#### 3.2.5. Speed of action reprogramming

The results of the GLMM run on time between the peak of the partial burst and the re-initiation of the unimanual movement are depicted in Figure 6 and Table 2. There was a significant main effect of experiment ꭓ^2^(1) = 48.74, *p* < 0.001, whereby action reprogramming was faster in the ARIignore condition and a significant main effect of trial type ꭓ^2^(1) = 9.86, *p* = 0.002, whereby action reprogramming was faster in stop trials than ignore trials. The also revealed a significant interaction between condition and trial type ꭓ^2^(1) = 4.41, *p* = 0.03. Tests of simple main effects revealed that this was due to action reprogramming to ignores being slower than that to stops in the ARIignore condition (*z* = 3.43, *p* < 0.001), but not in the SSTIignore condition (*z* = 1.46, *p* = 0.141). The covariate of age was significant ꭓ^2^(1) = 18.24, *p* < 0.001, with greater age being associated with a longer duration of reprogramming β = 1.009 (95%CIs 1.006, 1.013).

**Figure 6.**
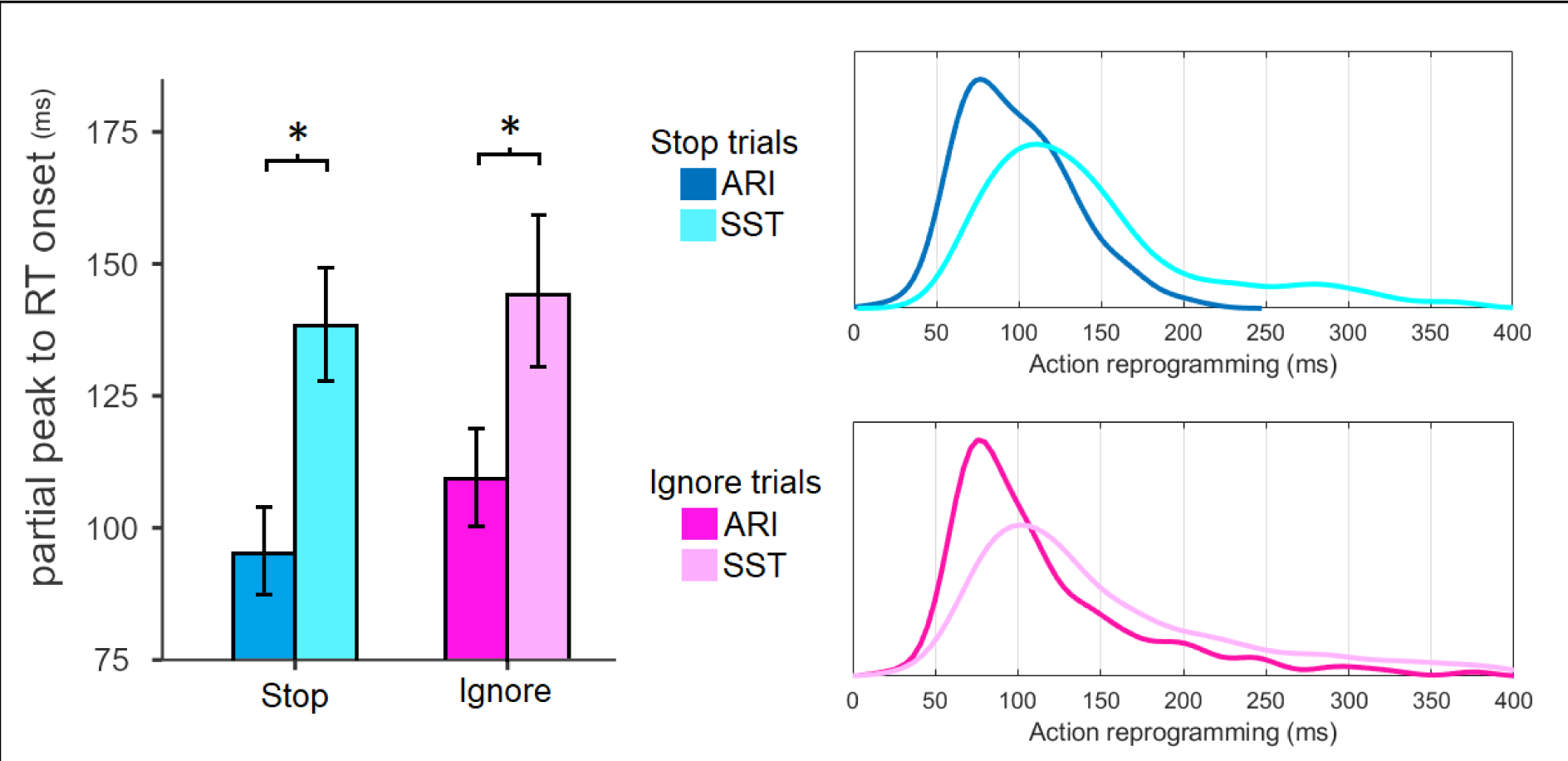
Speed of action reprogramming (time from the peak of the partial burst to the onset of the RT-generating burst in milliseconds) in stop and ignore trials in the ARIignore and SSTignore conditions (in the figure, referred to simply as ARI and SST, to aid interpretation). Note that distributions represent *pooled* data, while our GLMM included random intercepts for participants and random slopes for condition and trial type. The y-axis label is omitted here as the trial densities are relative to each category, to allow for clearer visual comparisons * = *p* < 0.001.

## 4. Discussion

The current study used behavioural and physiological measures to directly compare stopping behaviour between stop signal tasks (SSTs) and anticipated response inhibition (ARI) tasks where the tasks required both stimulus- and motor-selective stopping. To our knowledge, no prior research has directly compared EMG-based measures of reactive inhibition (CancelTime) on a within-subjects basis between these two tasks. Here, we report no evidence of differences in reactive inhibition between tasks, suggesting the tasks rely upon the same underlying inhibitory mechanisms. We also use EMG-derived measures to provide novel insights into the speed of action reprogramming in both tasks, concluding that the shorter stopping delays observed in the ARI task are a result of faster action reprogramming, rather than faster action cancellation.

### 4.1. Reactive inhibition was not significantly different across task contexts

Our model comparing CancelTime across the different stopping tasks (ARI and SST; with and without ignore trials) revealed no significant differences between SSTs and ARI tasks. A follow-up Bayesian ANOVA (JASP Team 2018, Morey & Rounder 2018, Rounder et al., 2012) confirmed strong evidence for the null hypothesis (BF_01_ = 15.99; see supplementary analysis). This provides the first physiological (EMG-based; CancelTime) evidence that the speed of action cancellation does not vary between SST and ARI tasks. Past research using behavioural measures (SSRT) to investigate motor selective stopping have reached similar conclusions (Hall et al., 2022; Wadsley et al., 2023), though other research which featured simple stopping (i.e., the go and stop responses involved only one effector) have observed faster SSRTs in ARI tasks compared to SSTs (Leunissen et al., 2017). These differences could be a result of the unreliable nature of conventional methods of SSRT calculation in ARI tasks (Matzke et al. 2021) and selective stopping tasks (Bissett & Logan, 2014) which are not a simple race between a stop and go process, but also involve a third unimanual response (Gronau et al. 2023). Alternatively, it may be that when only one effector is engaged in action and subsequent cancellation (i.e., simple, non-selective stopping), the inhibition of anticipated movements is faster than that of reactive movements (as per Leunissen et al., 2017). Further research directly comparing CancelTime in simple and motor-selective SSTs and ARI tasks would clarify this.

The percentage of successful stop trials in which partial bursts were elicited (i.e., muscle activation was initiated but cancelled before resulting in an overt button press) was significantly greater in the SSTs than that in the ARI tasks (refer to Table 2 and section 3.2.1). For a partial burst to be detected, the initial action must first have reached the periphery before the inhibitory response then terminates (cancels) this action: If inhibition occurs too early, peripheral muscle activation will not have been initiated; there will be a successful stop with no partial burst. If inhibition is too late, stopping can be unsuccessful or successful, but in the latter case the partial burst will not be temporally distinguishable from the subsequent EMG resulting in the RT generating burst in the responding (non-stopping) hand, with EMG of the two processes merging (Gronau et al., 2023; Salomoni et al. 2023). Only between these two constraints will a partial burst be reliably detected and characterised as a cancelled response. The reason for the smaller number of partial bursts in the ARI tasks is not entirely clear, although the narrower distribution of go RTs (Figure 2) may contribute. To explore this further, a supplementary analysis was undertaken on EMG data from the responding hand in correct stop trials. This revealed that the average time from EMG onset to when a button press was registered (the period within which an action needs to be interrupted to result in a partial burst) was significantly shorter in the ARIignore condition compared to the SST, although this difference was not significant in the ARI condition. Such a theory, therefore, does not fully explain the dichotomy in the extent of partial bursts observed. It is also possible that the faster speed of action reprogramming in the ARI tasks (Figure 6), allows for a greater number of trials in which inhibition occurs at the cortical level before any motor command is evident at the level of the muscle. As a practical consideration, we note that if researchers are primarily seeking to calculate inhibition latency using EMG, SSTs are more likely to yield interpretable results due to the greater proportion of partial bursts.

Although we have included calculations of SSRT in the SSTs, we note that these should be interpreted with caution due to well documented limitations with traditional SSRT calculation methods (Bissett et al., 2019; Gulberti et al., 2014; Matzke et al., 2017; Verbruggen & Logan, 2015), especially in stimulus selective stopping contexts (Bissett & Logan, 2014).

### 4.2. Stopping delays are shorter in ARI tasks than SSTs

Stopping delays – i.e., the extent to which RTs in successful selective stop trials are slower than go RTs - were smaller in the ARI tasks than the SSTs (refer to sections 3.1.3 and 3.2.4). Visual inspection of the distributions of stopping delays demonstrate that these delays vary widely from no delay (relative to that participant’s mean go trial RTs) through to trials with very large delays. It should be some of this variability is likely driven by comparing individual selective stop RTs to a single (average) go RT value, when we know substantial variability exists in go RTs (figure 2). Nonetheless, the stopping delay was also dependent on SSD, with longer SSDs being associated with the longest delays. This is consistent with a recent model of selective stopping which posits that the stop signal triggers two simultaneous processes: a non-selective inhibitory response (affecting both hands) and a unimanual go response (for the non-cued hand; e.g., left hand in a stop-right trial; Gronau et al., 2023). It follows that a later stop signal results in a later starting point for the unimanual response from the responding hand, and thus a longer stopping delay (relative to the go stimulus). Overall, our analysis on action reprogramming suggests that shorter stopping delays in ARI tasks (c.f. SSTs) result from faster action reprogramming, following cancellation of the initial action.

During preparation of an anticipated movement there is a preparatory *release* of intracortical inhibition in the motor cortex, facilitating a rapid response (Davranche et al., 2007; Sinclair & Hammond, 2008, 2009, Hannah et al., 2018). Such modulations may also contribute to the faster action reprogramming in the ARI task, where the timing of the go response is predictable. Though we note that for this to be the case, the modulations to intracortical inhibition would need to persist – at least to some extent - following the presentation of a stop signal (which has been associated with an *increase* of intracortical inhibition; Coxon et al., 2006; MacDonald et al., 2014; Stinear et al., 2009).

### 4.3. Stopping delays are not significantly different between stop and ignore trials when partial bursts occur

We used EMG to partition stop and ignore trials into those with partial bursts and those without. While previous papers have identified partial bursts only in successful stop trials (Raud et al., 2022), here we generalise this finding to both stop and ignore trials (refer to section 2.8.1). Moreover, we apply our algorithms to be able to detect partial bursts in the stopping hand as well as the continuing hand (Weber et al. 2023), consistent with models of global inhibition (MacDonald et al.2017; Gronau et al. 2023) and facilitating the action reprogramming analyses. The EMG profiles depicted in Figures 3 and 4 demonstrate the presence of inhibition in response to ignore cues in both the SSTignore and ARIignore conditions. Consistent with our previous findings (Weber et al., 2023), the current stimulus-selective SST revealed no significant difference in stopping delays between stop and ignore trials when trials with partial bursts were analysed. Here, we extend these findings, demonstrating this finding generalises across tasks (i.e., for both ARI tasks and SSTs). When only behavioural results are considered, stopping delays appear smaller in response to ignore trials than stop trials (Ko & Miller, 2013; Wadsley et al., 2023). Furthermore, failing to respond to the signal presented will result in an incorrect response in stop trials, whereas this trial will still be counted as correct in ignore trials, further reducing mean stopping delays to this trial type in the behavioural data. The current findings therefore support the interpretation that this behavioural finding is the result of averaging over a smaller *proportion* of trials where global inhibition occurred in response to ignore trials, rather than true differences in the magnitude of the effect. The EMG analysis presented here supports this argument by showing significant differences between stop and ignore trials only in the absence of partial responses, but no differences in trials where participants responded to the signal presented, inhibiting their initial bimanual response.

### 4.4. Reactive inhibition and action reprogramming, but not proactive inhibition, were influenced by age

Consistent with previous research, we observed a reduction in the speed of reactive inhibition with advancing age (Bloemendaal et al., 2016; Coxon et al., 2016; Hsieh & Lin, 2016; Kleerekooper et al., 2016; Sebastian et al., 2013; Smittenaar et al., 2015; Weber et al., 2023). However, in contrast with previous research (Bedard et al., 2010; Sebastian et al., 2013; van de Laar et al., 2011; Weber et al., 2023) no significant effects of age were observed with regards to proactive slowing ^2^. Given that the oldest participant in the current study was 55 years old, this result may suggest that changes to proactive slowing are more prominent in later senescence, rather than increasing incrementally throughout adulthood. This notion is supported by work from Bedard et al., 2010, which featured an age range of 6-82 years and observed the largest jump in proactive slowing in the 60-82 years age-group. Alternatively, it is possible that our post-trial feedback mitigated the extent of proactive slowing, such that it was kept within a limited range for all participants, regardless of age.

A novel finding of the current study is that action reprogramming during both anticipated (ARI) and reactive (SST) versions of selective stopping tasks slows with advancing age, alongside the aforementioned slower reactive inhibition. This finding contributes to the evidence of decreases in motor control which occurs in later life, and directly indicates that motor control is particularly impaired in complex stopping situations, requiring the coordination of numerous effectors and action reprogramming.

### 4.5. Conclusion

Our results demonstrate that the speed of the inhibitory response (as indexed by EMG) is not significantly influenced by whether the enacted movement is reactive (as in the SST) or anticipated (as in the ARI task). However, the selective inhibition of anticipated movements is more efficient as a result of faster action reprogramming. This novel finding helps to explain the stopping delays observed in ARI tasks, relative to SSTs. Finally, the latency of the inhibitory response, and the speed of action reprogramming are influenced by age, with older age being associated with both slower inhibition and slower action reprogramming in both anticipated and reactive response tasks.

## Author contributions

**Simon Weber:** conceptualisation, methodology, software, writing – original draft, formal analysis, investigation. **Sauro Salomoni:** conceptualisation, methodology, software, formal analysis, writing – review & editing. **Mark Hinder:** conceptualisation, resources, writing – review & editing, supervision, project administration, funding acquisition.

## Supplementary Material

### Time from EMG onset to RT-generating burst

To explore potential explanations for the smaller number of partial bursts in the ARI tasks compared to the SSTs (refer to Table 2), a supplementary analysis examining the time between the onset of EMG activity and the time of the registered button press was undertaken. A generalized linear mixed model was run with a gamma distribution and a log link function. Fixed effects featured a single condition with 4 levels (ARI, ARIignore, SST, SSTignore) The random effects structure included participant intercepts and slopes for condition. The analysis revealed a main effect of condition ꭓ^2^(3) = 16.97, *p* < 0.001. Bonferoni corrected post-hoc tests revealed EMG-onset to RT was shorter in the ARIignore condition than the SST condition (*z* = 3.05, *p* = 0.014, and the SSTignore condition (*z* = 4.03, *p* < 0.001). However, EMG-onset to RT was not shorter in the ARI condition than the SST condition (z = 1.44, p = 0.91) or the SSTignore condition (*z* = 2.41, *p* = 0.09). Estimated marginal means are presented in Table S1.

**Table S1.**
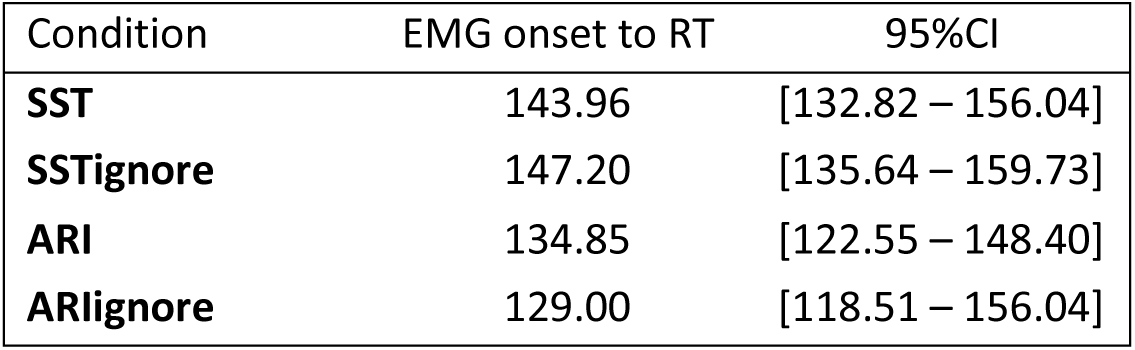

### Bayesian Analysis to confirm no difference in CancelTime between conditions

Following our initial GLMM run on CancelTime we decided to include a Bayesian analysis, to provide evidence that the result was due to our analysis being under-powered. The ANOVA used a single condition with four levels (ARI, ARIignore, SST, SSTignore).

Model Comparison – CancelTime

Model Comparison - CancelTime
| Models | P(M) | P(M\|data) | BF<sub>M</sub> | BF<sub>01</sub> | error % |
| --- | --- | --- | --- | --- | --- |
| Null model | 0.5000 | 0.9412 | 15.9959 | 1.0000 |  |
| Condition | 0.5000 | 0.0588 | 0.0625 | 15.9959 | 0.0221 |

## Footnotes

1 While correct ignore trials have an RT for both the cued and non-cued hand, we used the RT from the non-cued hand to ensure stopping delays were measured the same way in stop and ignore trials.

2> We note that Smittenaar et al., (2015) also observe no changes in proactive *control* in older adults, though this measure was related to the use of cues to inform stopping, and is distinct from typical calculations of proactive slowing.

